# Simulation-based approaches to characterize metagenome coverage as a function of sequencing effort and microbial community structure

**DOI:** 10.1101/356840

**Authors:** Taylor Royalty, Andrew D. Steen

## Abstract

We applied simulation-based approaches to characterize how microbial community structure influences the amount of sequencing effort to reconstruct metagenomes that are assembled from short read sequences. An initial analysis evaluated the quantity, completion, and contamination of complete-metagenome-assembled genome (complete-MAG) equivalents, a bioinformatic-pipeline normalized metric for MAG quantity, as a function of sequencing effort, on four preexisting sequence read datasets taken from a maize soil, an estuarine sediment, the surface ocean, and the human gut. These datasets were subsampled to varying degrees of completeness in order to simulate the effect of sequencing effort on MAG retrieval. Modeling suggested that sequencing efforts beyond what is typical in published experiments (1 to 10 Gbp) would generate diminishing returns in terms of MAG binning. A second analysis explored the theoretical relationship between sequencing effort and the proportion of available metagenomic DNA sequenced during a sequencing experiment as a function of community richness, evenness, and genome size. Simulations from this analysis demonstrated that while community richness and evenness influenced the amount of sequencing required to sequence a community metagenome to exhaustion, the effort necessary to sequence an individual genome to a target fraction of exhaustion was only dependent on the relative abundance of the corresponding organism and its genome size. A software tool, GRASE, was created to assist investigators further explore this relationship. Re-evaluation of the relationship between sequencing effort and binning success in the context of the relative abundance of genomes, as opposed to base pairs, provides a framework to design sequencing experiments based on the relative abundance of microbes in an environment rather than arbitrary levels of sequencing effort.

## Background

The reconstruction of high-accuracy short read sequences into metagenome-assembled genomes (MAGs) is a recent approach to characterize microbial metabolisms within complex communities [1]. The recent creation of ~8,000 MAGs from largely uncultured organisms across the tree of life [2], the spatial characterization of microbial metabolisms and ecology across Earth’s oceans [3], and the characterization of the potential impact that fermentation-based microbial metabolisms have on biogeochemical cycling in subsurface sediment environments [4] provide a few examples of how MAGs have helped to elucidate the relationships between microbial ecology, microbial metabolisms, and biogeochemistry.

Sampling environmental microbial DNA involves selecting a target environment, sequencing effort, bioinformatic pipeline software and parameters, metabolism characterization software (i.e., for gene identification and similarity searches) and databases (Fig. 1). At present, there is little information to guide how much sequencing is appropriate to achieve scientific goals in such experiments [5]. This gap in knowledge is partly attributed to the unknown structure of target microbial communities. Further complications arise in that the accuracy and efficiency of bioinformatic pipelines are often difficult to characterize, and thus obscure the relationship between sequencing effort and MAG retrieval. Recent estimates compiled by Quince et al. [5] suggest that typical metagenomic shotgun sequencing experiments usually sequence between 1 Gbp and 10 Gbp. Researchers require more precise guidance to select an appropriate shotgun sequencing effort in order to maximize information and minimize cost for their specific experimental question.

**Fig. 1.**
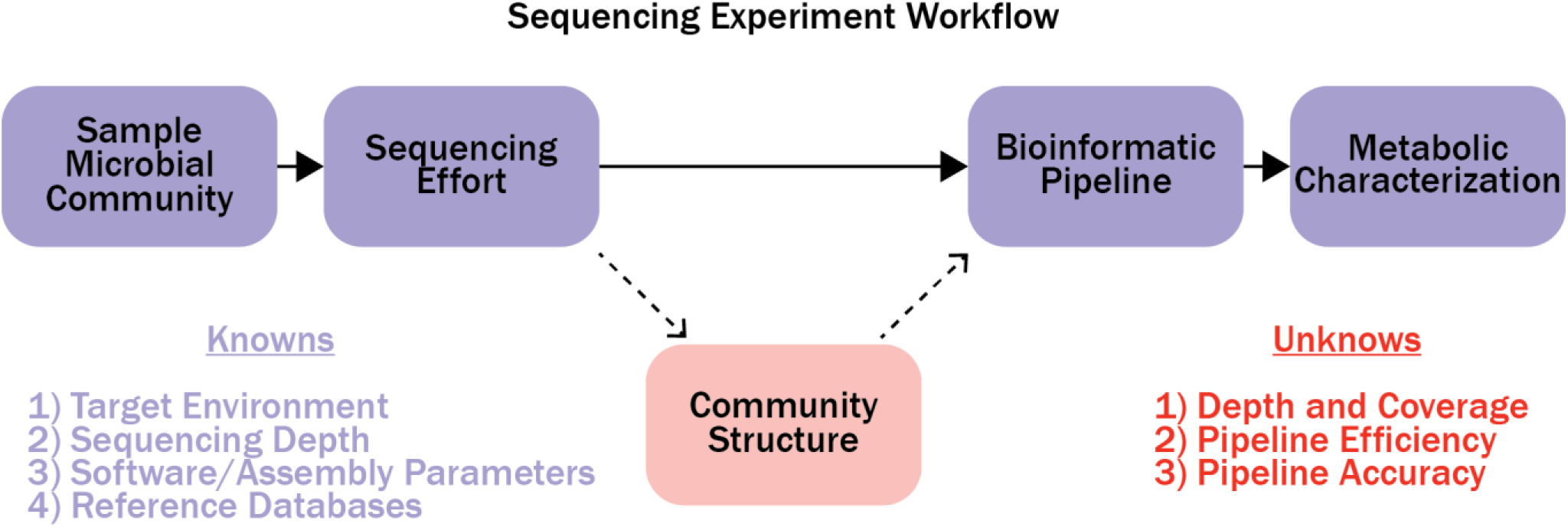
A flow diagram illustrating the workflow for sequencing experiments.

Illumina sequencing technology is currently the most popular platform to generate metagenomic shotgun sequences [5]. Previous investigators established theoretical relationships between contig formation rate [6] and single genome coverage [7] as a function of short read sequencing effort. On the community level, heuristic approaches have been proposed for evaluating community-level coverage to increases in sequencing effort. For example, it has been proposed to utilize short read redundancies as a function of sequencing effort to estimate community level coverage [8]. Without *a priori* knowledge of the microbial community structure, practical application of these methods to estimate MAG retrieval as a function of sequencing effort is hindered. This is particularly true as heuristic methods will only be sensitive towards highly abundant genomes.

Here we present two distinct analyses which constrain the relationship between the quantity of Illumina metagenomic shotgun sequences and the community-level sequence coverage. First, we performed *in silico* experiments to simulate the effect of sequencing effort on retrieved MAG properties for Illumina sequence datasets. For these experiments, we propose a new metric, complete-MAG equivalents, which measures community coverage in response to sequencing effort and is insensitive to splitting versus joining bins in our bioinformatic pipeline. Second, we applied a theoretical model and numerical simulations to estimate the minimum sequencing effort needed to sequence a metagenome to a target fraction of exhaustion. Our theoretical model is unique compared to previous models [6,7] in that we characterize sequencing effort in the context of all physically-distinct DNA fragments in the metagenome. This is important, as complete genome coverage does not necessarily translate into accurate read assembly due to interspecies homologous DNA and intraspecies microdiversity [9]. Coupling results from the two analyses provides a framework for investigators to define sequencing experiments in the context of selecting a rarity and fraction of exhaustion for a desired target genome when sequencing a community. The patterns presented here can be used to guide sequencing effort decisions in future sequencing projects when MAG reconstruction is a primary goal.

## Results

### MAG RECONSTRUCTION AS A FUNCTION OF SEQUENCING EFFORT IN EXISTING METAGENOMIC DATASETS

The number of ‘complete-MAG equivalents’ (equation 1) as a function of high-quality bases empirically fit the Gompertz equation (equation 2; Fig. 2b; parameters in Table 1; Additional file 1). Complete-MAG equivalents represent the sum of MAGs retrieved, weighted by each MAG’s checkM completeness score [10], and divided by 100% (i.e., a complete MAG). This metric is designed to highlight the total number of marker genes, used to determine MAG coverage, in an environment while maintaining the biological relevance of MAGs. Complete-MAG equivalents reduces sensitivity to whether bioinformatic pipelines tend to lump contigs into fewer, more-complete MAGs versus split them into fewer, less-complete MAGs.

**Table 1.** Estimates of fit coefficients for the Gompertz equation (equation 3) for the complete-MAG equivalents as a function of sequencing depth in published datasets from ocean surface water, estuarine sediment, maize soil, and the human gut. *p* values for all coefficients were <<0.05.

| Environment | $A$ (±SE) | $\mu$ (±SE) | $\lambda$ (±SE) | Yield* |
| --- | --- | --- | --- | --- |
| Ocean Surface Water | 97.67 (4.15) | 5.84 (0.31) | 1.16 (0.33) | 0.88 |
| Estuary Sediment | 26.25 (2.11) | 1.63 (0.06) | 3.70 (0.24) | 0.86 |
| Maize Soil | 43.65 (1.98) | 1.43 (0.07) | 6.13 (0.60) | 0.70 |
| Human Gut | 17.49 (1.02) | 5.01 (0.46) | 0.67 (0.14) | 0.90 |

\*Equation 3

**Fig. 2.**
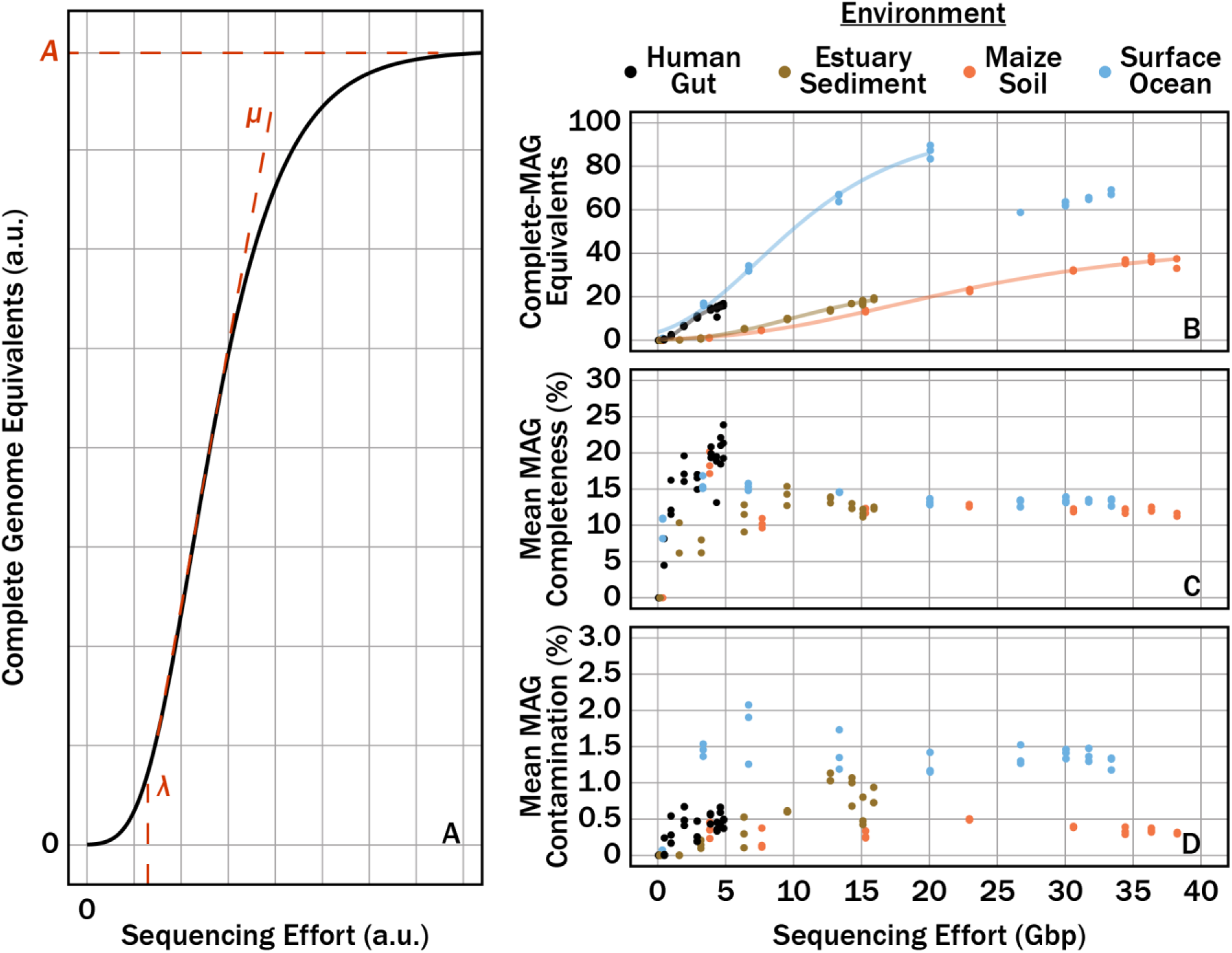
The influence that the parameters *A*, *μ*, and *λ* had on the Gompertz equation (a). The property of the Gompertz equation that each parameter influences is colored red. Complete-MAG equivalents (in arbitrary units) (b), mean MAG completeness (c), mean MAG contamination (d) as a function of simulated sequencing effort (Gbp) for sequence datasets of the human gut, maize soil, estuarian sediment, and surface ocean microbiomes, using the pipeline described in the methods section. Translucent lines in (b) correspond to nonlinear least squares fits of the Gompertz equation to the respective environmental dataset.

For each environment, the number of complete-MAG equivalents as a function of simulated sequencing effort fit the Gompertz equation (equation 2). Here, *A*, *μ,* and *λ* correspond to the maximum number of complete-MAG equivalents assembled with the pipeline, the maximum rate which complete-MAG equivalents form with more sequencing, and the “lag bases,” or the bases which must be sequenced prior to a sufficient number of sequence reads existing to generate overlap and form contigs [6]. The predicted maximum complete-MAG equivalents varied from ~17 in the gut community to ~97 in the surface ocean. The predicted maximum rate that complete-MAG equivalents increased varied from ~1.4 to ~5.8. Lastly, the minimum threshold of sequencing necessary prior to seeing complete-MAG equivalents varied from ~0.6 to ~6.7. The *Tara Oceans* dataset, where complete-MAG equivalents decreased at sequencing effort >20 Gbp, was an exception. For the estuary, maize, and human gut datasets, the complete-MAG equivalents yield began to asymptote with increasing sequencing efforts. The *Tara Oceans* dataset followed a similar pattern at <25 Gbp. However, when the number of sequenced bases was >25Gbp, the number of complete-MAG equivalents decreased and became insensitive to sequencing effort. The total number of MAGs retrieved was considerably higher than the number of complete-MAG equivalents.

Mean MAG completeness also increased towards an asymptote with increasing sequencing effort (Fig. 2c). Completeness was highest for the human gut dataset, with a maximum of 23.9%, and increased continuously as sequencing effort increased. The mean MAG completeness reached an asymptote of ~10-15% for the other three datasets with sequenced bases >10 Gbp. Note that when >10 Gbp were sequenced, the number of complete-MAG equivalents created still increased as new sequences were added. For all datasets, mean MAG contamination was <2% (Fig. 2d) and did not depend strongly on sequencing effort.

### SIMULATION EXPERIMENTS

Using equation 9, we calculated the number of *k*-length sequence reads required to sequence all physically-distinct DNA sequences of length, *k* (*k*-mers), in four hypothetical metagenomes, a perfectly even, moderately uneven, highly uneven, and a lognormally-distributed structure (Fig. 3a-d). Here, we define physically-distinct DNA as all possible *k*-mer combinations comprising a genome. Note that *k*-mers within homologous regions are mathematically treated independent. The log-transformed expectation value of sequencing effort (number of sequences) required to fully sequence genomes of the hypothetical communities was linear with respect to log-transformed size of the metagenome (i.e., physically-distinct *k*-mers in the population, or unique base pairs in a metagenome); this suggests a power-law relationship between metagenome size and expectation value of sequence reads required to sequence the metagenome to exhaustion (Fig. 3e). For all community structures, the slope of the relationship between sequencing effort (log-transformed sequenced reads) and metagenomic richness (log-transformed physically-distinct number of sequenced reads) was within 1% of 1.06. The structure of the population strongly influenced the number of reads required such that more-even community structures required far fewer reads than less-even structures.

**Fig. 3.**
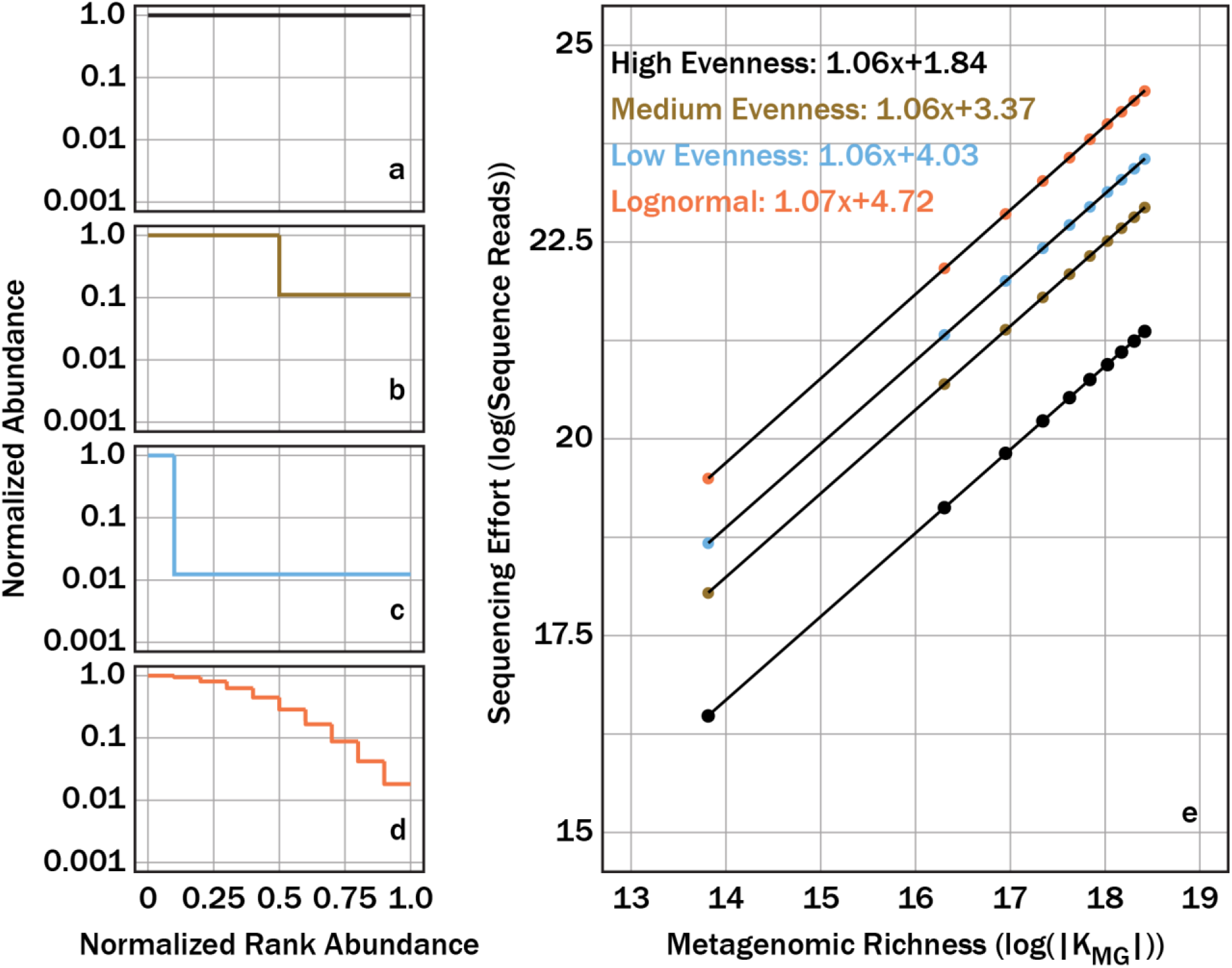
Sequencing effort, with units of log of sequence reads, required to fully sequence four different community structures, one with relatively high community evenness (a), relatively moderate community evenness (b), relatively low community evenness (c), and one with a lognormal community structure (d), were predicted using linear regressions (e) and the log of |*K_MG_*| from equation 9 as a predictor.

As equation 9 only estimates the number of reads to sequence a metagenome to exhaustion, we used a numerical simulation to estimate the number of *k*-sized reads to sequence a metagenome to a target fraction of exhaustion. The numerical simulation and equation agreed on the expected sequences reads to sequence 100% of a given metagenome (Fig. 4); this supported the use of this simulation. The log-transformation of both total physically-distinct *k-*sized reads (|*K*_MG_| in equation 8) and sequenced reads showed a linear response for all target fractions and all community structures. The amount of sequences required to achieve a given target of physically-distinct *k-*sized reads was variable for the different communities shown in Fig. 3a. For instance, the lognormally-distributed community, when compared to other communities, required the most sequencing effort to sequence a metagenome to exhaustion but required similar sequencing effort to sequence a metagenome to a target fraction of 50%.

**Fig. 4.**
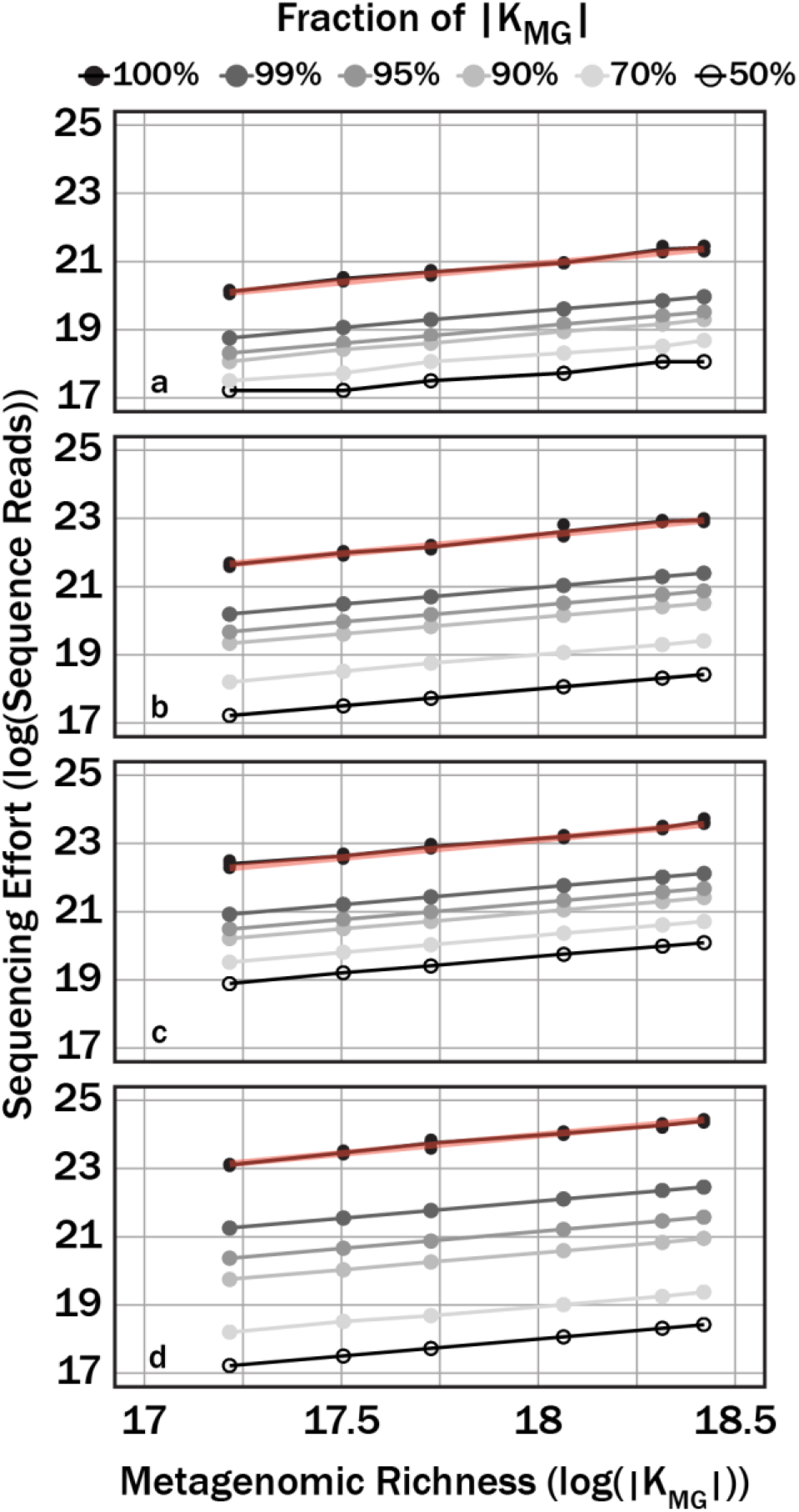
Sequencing effort, with units of log sequence reads, necessary to reach variable target sequencing depths (colors) for four different community structures, one with relatively high community evenness (a), relatively moderate community evenness (b), relatively low community evenness (c), and one with a lognormal community structure (d). Red translucent lines correspond with linear regression curves for the respective community in Fig. 3e.

We applied the simulation to demonstrate the effect of community evenness on the number of reads required to sequence a community to a target fraction of completion. These communities ranged from perfectly even (*a*=0, equation 11) to more uneven (*a*=0.02, Fig. 5a). Evenness was quantified using the Pielou evenness index, which expresses Shannon diversity relative to the diversity of a perfectly even community [11]. The sequencing effort required to characterize genomes depended strongly on both the evenness and the target fraction of completion (Fig. 5b). Again, less-even communities required more sequence reads than more-even communities. The strength of this relationship also depended on the target fraction of completion. A community with Pielou evenness of 0.97 required 3 orders of magnitude more sequence reads to sequence a metagenome to a target fraction of exhaustion than a perfectly even community while the same community only required about 42% more reads to sequence 50% of the metagenome.

**Fig. 5.**
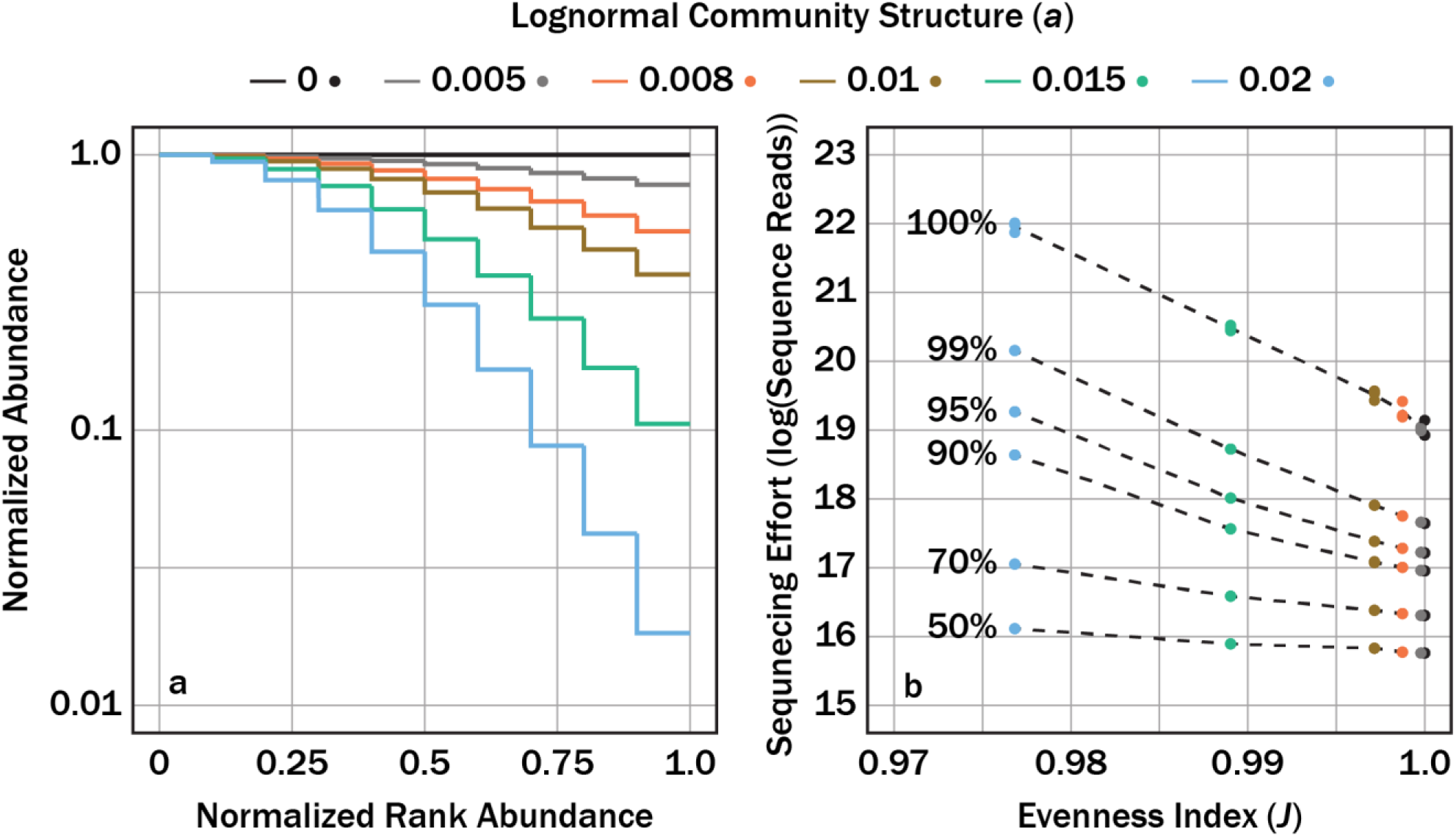
Numerical sequencing simulations applied to 6 hypothetical communities with different lognormal distributions that were defined by the parameter, *a*, from equation 10 (a). The sequencing effort, with units of log of sequence reads, necessary to sequence a target fraction of |*K_MG_*| (dashed contours) as a function of the Pielou evenness index, *J*, for a given lognormal community structure (b).

The minimum sequencing effort required to sequence a microbe genome given a combination of target fraction, genome size, and fraction of the metagenome community was modeled with a generalized additive model (GAM; Fig. 6). The GAM shows expected sequencing effort required for microbial genome sizes 0.5, 2, 5, 10, and 20 Mbp, target genome completeness fractions from 0.5 to 1.0, and genome relative abundances from 1 to 0.0001. The smooth dimensions for target fraction, genome size, and fraction of the metagenome community were 50, 6, and 29, respectively. To normalize for different sequence read length, sequence reads were converted to bases and ranged from 1×10^7^ to 1×10^15^ total bases. More bases were required to sequence microorganisms 1) when the genome was relatively rarer in the community, 2) to achieve better coverage of the genome, and 3) when average genome sizes were larger.

**Fig. 6.**
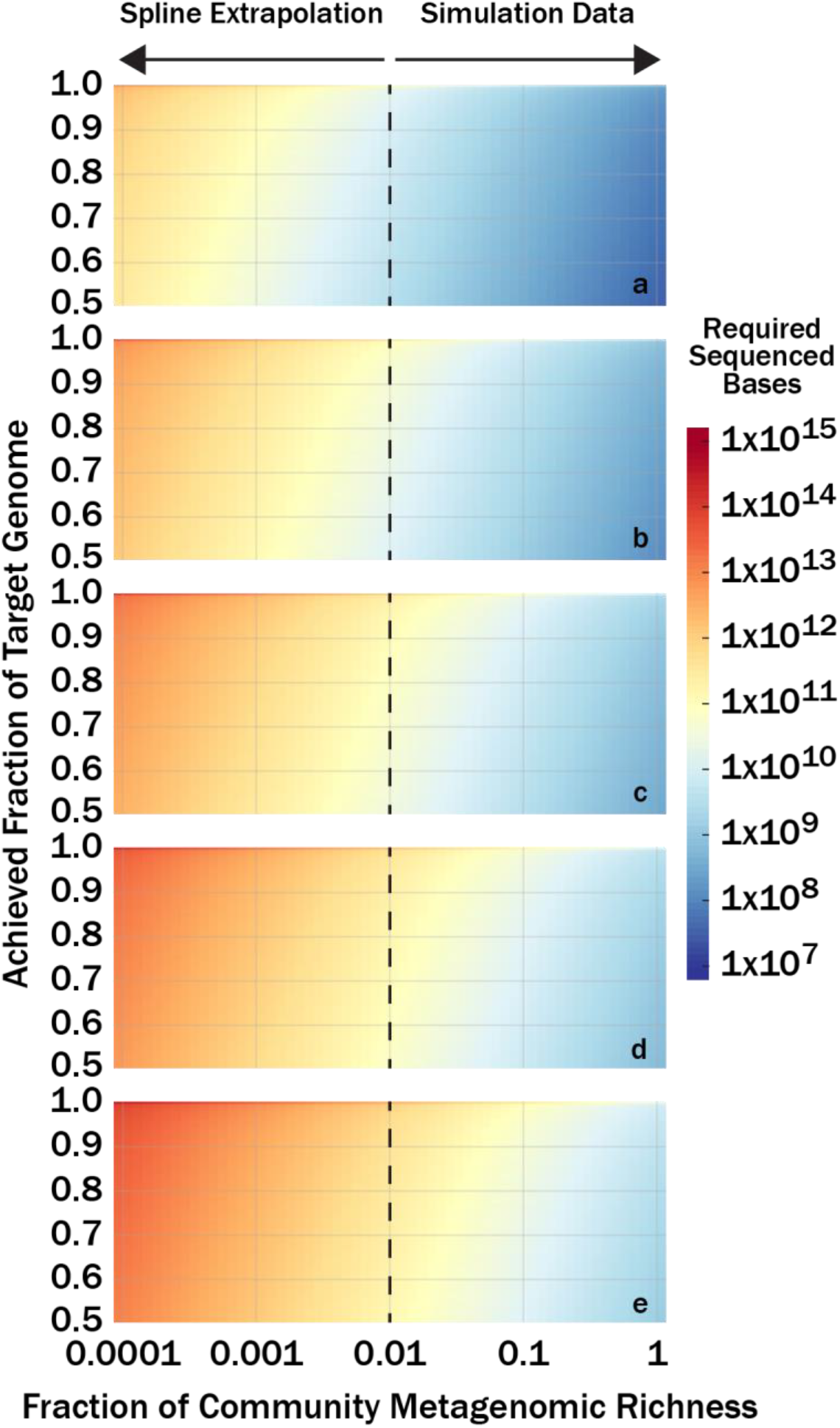
Numerical sequencing simulations show the number of bases (color bar) required to sequence a target fraction of a genome as a function of that genome’s relative abundance in the community metagenome. Genomes evaluated were 0.5×10^6^ (a), 2×10^6^ (b), 5×10^6^ (c), 10×10^6^ (d), and 20×10^6^ (e) base pairs long.

Using the relationship shown in Fig. 6, we can convert sequencing effort to an abundance cutoff. Fig. 7 shows complete-MAG equivalents retrieved for a genome of a given level of abundance that is sequenced to a target depth for the human gut, maize soil, estuarine sediment, and the surface ocean microbiomes analyzed earlier. Correlation coefficients for the regressions used to relate log-transformed sequencing effort (base pairs) to genome relative abundance were *R*=1 for all three genome sizes evaluated (1 Mbp, 5 Mbp, and 20 Mbp). Evaluation of 1 Mbp and 20 Mbp define the range of uncertainty in predicting complete-MAG equivalents as the true size of genomes is unknown. Unlike the asymptotic response of complete-MAG equivalents to sequencing effort (Fig. 2b), complete-MAG equivalents increase exponentially as the abundance cutoff decreases (note that the abundance cutoff is on a log-scale). The target genome completeness fraction was held at a constant of 0.5 for all regressions and the sensitivity of complete-MAG equivalents will change with respect to the genome abundance cutoff with different values of target genome completeness fraction.

**Fig. 7.**
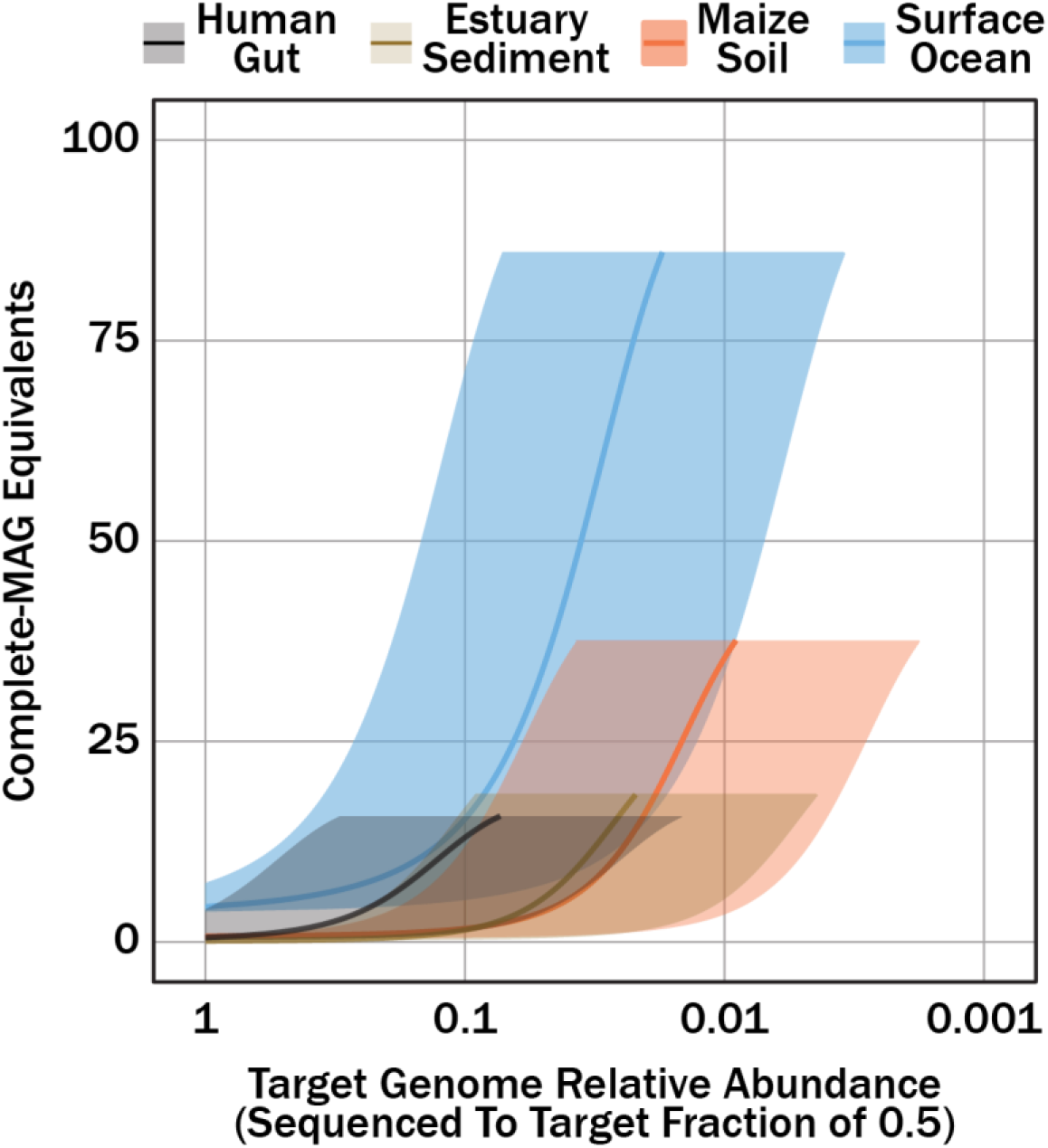
Complete-MAG equivalents as a function of a target genome’s relative abundance when sequenced to a target fraction (0.5) for human gut, maize soil, estuarian sediment, and surface ocean sequence datasets. Sequencing effort was converted to genome relative abundance sequenced to target fraction using the GAM presented in Fig. 6. The translucent shaded areas correspond to uncertainty from the target genome size (1 Mbp, or the lower-bound, to 20 Mbp, or the upper-bound) while the solid lines correspond to genome sizes 5 Mbp.

## Discussion

We sought to establish evidence-based guidelines for selecting a sequencing effort during shotgun metagenomic sequencing experiments. Random subsamples of existing short read datasets (individually sampled, assembled, and binned) simulated the effect of creating MAGs from datasets of different sizes and environments. The datasets analyzed here are representative of both the sequencing effort (1 to 10 Gbp) [5] and the types of target environments that microbial ecologists often investigate [12]. A variety of software is available for all steps of MAG creation pipelines, and the quantity/quality of MAGs will depend on software selection, software configuration, and sequenced environment (Fig.1) [5]. Furthermore, it is best-practice to manually curate algorithmically-created MAG bins [13]. We do not argue that the pipeline used here is objectively optimal for generating high-quality (high completeness and low contamination) MAGs. Rather, our pipeline was configured to minimize contamination (Fig. 2D) associated with retrieved MAGs at the expense of reduced completeness. Thus, MAG quantity was not directly reported but expressed as complete-MAG equivalents (equation 1). Reporting complete-MAG equivalents reduces bias associated with binning software.

As sequencing effort increased, there was an initial lag in complete-MAG equivalents followed by a rapid increase in complete-MAG equivalents, and then diminishing returns at higher sequencing efforts. Previous investigators modeled the response of 16S RNA gene [14–16], Hill’s number diversity [17], taxon-resolved abundance [18], and gene abundance [18] as a function of sequencing effort with rarefaction curves, or collector’s curves. The complete-MAG equivalents as a function of sequencing effort did not match a traditional collector’s curve, which lacks an initial lag. Rather, the data appear sigmoidal. We modeled the data using the Gompertz function (equation 9), because its parameters can be interpreted in terms of quantities that are familiar from microbial growth curves (lag time, growth rate, and maximum density) [19]. The three Gompertz parameters, *A*, *μ*, and *λ* are interpreted as the maximum possible number of complete-MAG equivalents at infinite sequencing effort, maximum rate that complete-MAG equivalents increased with increasing sequencing effort, and a minimum sequencing threshold prior to rapid complete-MAG equivalents retrieval (Fig. 2a), respectively. Consequently, the Gompertz equation fit parameters may provide means for inter-comparisons between different metagenomic datasets. We want to emphasize that the datasets analyzed here do not necessarily reflect the generalized characteristics of their parent communities. More so, these communities were analyzed as their community properties (richness, abundances, and phylogeny) are generally different [4, 20–22].

The fits to the Gompertz function illustrate that there is an optimal sequencing effort for MAG creation efforts corresponding to the upper shoulder of the Gompertz curve (“late log phase”). When sequencing effort is too close to *λ*, MAGs bin poorly; when sequencing effort is too great, the number and quality of MAGs per unit sequencing effort (and therefore cost) is low. We speculate that our choice of pipeline, and specifically the fact that we discarded contigs <3kb, caused poor performance at higher sequencing effort for the *Tara Oceans* dataset. Species-level microdiversity and interspecies homologous DNA can cause “bubbles” which impair assembly in larger data sets [9,23]. Improved assembly would likely have yielded more complete-MAG equivalents for our assembly of the largest subsets of the *Tara Oceans* data.

As mean MAG completeness converged to an asymptote well below 100% (Fig. 2b), complete-MAG equivalent yields, or the ratio of the maximum observed complete-MAG equivalents to the theoretical maximum complete-MAG equivalents (*A*) (Table 1) were close to 100%. The asymptote in mean MAG completeness (Fig. 2c) and negligible values of mean MAG contamination (Fig. 2d) demonstrates that the retrieved complete-MAG equivalents at higher sequencing efforts is not attributed to artifacts of assembly. If the signal in complete-MAG equivalents was attributed to assembly of false positive marker genes, then either 1) the mean MAG completeness would decrease with the addition of false positive marker genes which fail to cluster (i.e., more MAGs with low completeness) or 2) false positive signal would cluster with preexisting MAGs leading to an increase in mean MAG contamination. Neither of these scenarios occurred. The fact that mean MAG completeness was at an asymptote <<100% before complete-MAG equivalents began converging suggests more sequenceable DNA exists in these communities. Thus, we asked how much sequencing was necessary to sequence a community to exhaustion.

The theoretical probability model (equation 9) demonstrated that the sequencing effort to sequence a metagenome to exhaustion was predictable, regardless of community structure (Fig. 3). Furthermore, a simplistic characterization community complexity demonstrates an inherent truth, less complex communities require less sequencing to sequence all available DNA. This corroborates our observations where less diverse communities (i.e., human gut) relative to the environmental communities had greater complete-MAG response to sequencing effort (Fig. 2b). A limitation to equation 9 is predicting sequencing effort to sequence a metagenome to a fraction of exhaustion. This is particularly important as 100% genome coverage is not necessary to obtain robust assemblages and MAGs. As such, a numerical simulation was used for predicting the sequencing effort necessary to sequence an individual metagenome to a fraction of exhaustion. The numerical simulation agreed with the theoretical model when predicting the sequencing effort to sequence a metagenome to exhaustion (Fig. 4). A last analysis explored the semi-quantitative relationship between community evenness and richness and the necessary sequencing effort to achieve a target fraction of exhaustion for a metagenome (Fig. 5).

In practice, information about a target community structure may not be available for estimating sequencing effort. The GAM built here predicts the minimum number of sequences necessary to sequence a given fraction of a target genome as a function of average genome size and the relative abundance of the target genome in the community (Fig. 6). Even without knowledge of a target community’s structure, the GAM provides a useful constraint for designing whole-genome shotgun metagenomic sequencing experiments. Note that the genome sizes selected for this analysis correspond with typical sizes reported for prokaryotic genomes [12]. Virus and eukaryotic genomes were not considered, as the primary focus of this study is prokaryotic systems. Contamination from eukaryotic and virus genomes is usually accounted for using size-fractionation. Nonetheless, the underlying theory applied here should be applicable to these other genomic systems.

Metagenomic shotgun sequencing experimental designs should be rationally designed such that sequencing effort is selected to capture a desired fraction of a target microbial genome. Investigators should be cognizant of the rarest microbial genome desired to be characterized as well as the degree of characterization of that microbial metagenome when designing a sequencing experiment. To that end, we have built a tool, `Genome Relative Abundance to Sequencing Effort (GRASE)`, to report estimated sequencing effort required to capture a defined fraction of a genome as a function of the relative abundance of the corresponding microorganism in the community and average genome size. This R-based GUI app can be accessed online at http://adsteen.shinyapps.io/grase and is archived at http://github.com/adsteen/GRASE, from which it can be downloaded and run locally.

When the sequence read datasets analyzed here (Table 2; Fig. 2) are re-evaluated in the context of the relative abundance of a microbial metagenome (*g*_MG_) sequenced to a target fraction (0.5), complete-MAG equivalents increase appreciably in response to minor increases in deeper characterization of the community metagenome (Fig. 7). This observation contrasts the complete-MAG equivalents response to sequencing effort (in base pairs), where substantial increases in sequencing effort (by contemporary standards) leads to diminishing returns in complete-MAG equivalents. It is important to note that abundance cutoff and sequencing effort are interchangeable; however, complete-MAG equivalents response to changes in the respective predictor (i.e., base pairs versus abundance cutoff) alters the optics of the collector’s curve. Modest increases in sequencing effort contribute minor amounts to extending abundance cutoff. A substantial amount of genomic and metabolic data can be gained from targeting rarer microbes (metagenomic abundances <0.005), with the caveat that whole-genome shotgun sequencing technology (as well as computational power) requires significant increases in either the number or length of reads generated per run.

**Table 2.** Summary of sequence datasets analyzed with the MAG pipeline.

| Environment | NCBI SRA Accession | Sequencing Platform | Total Reads* | High Quality Bases* | General Notes | Citation |
| --- | --- | --- | --- | --- | --- | --- |
| Ocean Surface Water | ERR599029 | Illumina HiSeq 2000 | $3.372 \times 10^8$ | $3.340 \times 10^{10}$ | Caribbean Sea (5 mbsl) | [20] |
| Estuary Sediment | SRR5248164 | Illumina HiSeq 2000 | $1.137 \times 10^8$ | $1.589 \times 10^{10}$ | Sulfate Zone (8-10 cmbsf) | [4] |
| Maize Soil | SRR351473 | Illumina HiSeq 2000 | $4.727 \times 10^8$ | $3.824 \times 10^{10}$ | Surface Soil | [21] |
| Human Gut | SRR5127631 | Illumina HiSeq 2000 | $5.095 \times 10^7$ | $4.847 \times 10^9$ | -- | [22] |

\* Combined forward and reverse paired-end reads.

## Materials and Methods

### SEQUENCE DATA SOURCES

All sequence data were downloaded from NCBI’s Sequence Read Archive (SRA) using the SRA Toolkit (fastq-dump –split-files) [24]. Exact duplicate reads for both forward and reverse reads were removed using PRINSEQ (-derep 1; v0.20.4) [25]. All sequencing datasets were limited to Illumina shotgun metagenomic paired-end reads. Four datasets were analyzed for this analysis. The first dataset was from oceanic surface water collected at 5m depth in the Caribbean Sea as a part of the *Tara Oceans* expedition [20]. The second dataset was from sediment from a depth of 8-10 cm below the surface (sulfate-rich zone) and collected at the White Oak River Estuary, Station H, North Carolina, USA [4]. The third dataset was collected from maize soil [21]. The last dataset was collected from human fecal samples and represented a human gut microbiome [22]. All datasets analyzed in this study are summarized in Table 1.

### MAG ASSEMBLY PIPELINE

The pipeline developed here followed similar pipelines described by other authors [3,26]. All sequence datasets were analyzed as follows. Trimmomatic (v0.36) [27] removed adapters and trimmed low-quality bases from individual reads. Read leading and trailing quality scores were required to be >3. The sliding window was set to 4 base pairs and filtered base pair windows with a mean score <15. Quality controlled reads were assembled into contigs using MEGAHIT (v1.1.2; --presets meta-large) [28]. Due to computational limitations, assembled contigs <3000 bp long were excluded from the analysis. Redundant contigs were removed using CD-HIT (v4.6.8; cd-hi-est -c 0.99 -n 10) [29]. Remaining contigs with high intra-contig similarity based on intra-contig sequence alignments (Minimus2 -D OVERLAP=100 MINID=95) were removed. The quality-controlled reads (i.e., after using Trimmomatic) were mapped to the remaining contigs using Bowtie 2 (v2.3.3) [30] to generate a coverage score for individual contigs.

Resultant contigs were iteratively clustered into MAGs using the unsupervised, clustering algorithm Binsanity (v0.2.6) [26]. Similar to Tully et al. [3], six initial clustering iterations were performed with the parameter, *preference* (-p), set to −10 (iteration 1), −5 (iteration 2), −3 (iteration 3-6). Between iterations, a refinement step (Binsanity-refine) was performed on the putative MAGs with constant *preference* (-p) of −25. The refined putative MAGs were evaluated for contamination and completeness using the software CheckM (v1.0.6) [10], which uses HMMER (v3.1) and Prodigal (v2.6.3) [31]. For this work, we used the recommended lineage-specific marker sets. The lineage-specific workflow defines marker genes as those appearing in >97% of a single lineage. Contigs associated with putative MAGs meeting one of the following criteria: 1) had a completeness > 90% and contamination < 10%, 2) had a completeness > 80% and contamination < 5%, or 3) had a completeness > 50% and contamination < 5% were treated as high-quality. All other MAGs were considered low-quality MAGs. MAGs defined as high-quality were not modified any further. Contigs associated with the high-quality MAGs were not used in the subsequent reclustering and refinement steps. The contigs associated with low-quality MAGs were pooled together and reclustered during the next iteration of Binsanity clustering. After the sixth iteration, the remaining MAGs which did not fall into one of the three categories underwent additional refinement using Binsanity-refine. During this step, MAGs were iteratively refined with *preference* set to −10 (iteration 1), −3 (iteration 2), and −1 (iteration 3). Between each refinement step, metrics of contamination and completeness were evaluated using CheckM. Again, MAGs meeting the criteria of a high-quality category described above were not further modified. The respective contigs associated with putative MAGs were not used in proceeding refinement steps. After the last iteration of refinement, all MAGs were reevaluated for completeness, contamination, and a taxonomic rank using CheckM. Completeness values for MAGs with a minimum resolved taxonomic rank of phylum were summed together. Complete-MAG equivalents, *g_eq_*, were defined as:

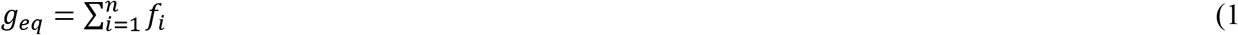

where *n* is the total number of retrieved MAGs and *fi* is the corresponding CheckM completeness (%).

Common practice when reporting MAGs involves discarding MAGs not meeting a criterion of completeness and contamination. In the context of characterizing MAG response to sequencing effort a MAG pipelines often fails at generating high-quality MAGs. One must discern whether this is a failure of the bioinformatic pipeline versus insufficient sequencing effort (Fig. 1). We argue that MAGs with low completeness still provide information regarding the community level coverage when generating collector’s curves. More so, a community has finite number of unique marker genes, and an infinite sequencing effort would result in sequencing all unique marker genes, regardless of whether or not these genes cluster to one another.

### SUBSAMPLING SEQUENCE READ DATASETS

The effect of decreased sequencing effort was simulated by subsampling the initial sequence read datasets described above. Downloaded sequence read datasets were randomly sampled at set fractions of 1%, 10%, 20%, 40%, 60%, 80%, 90%, 95%, and 100%. Each fraction was resampled, assembled, and binned in triplicate. Each triplicate assembly was binned independently using the MAG assembly pipeline described above.

### MODELING MAG RESPONSE TO SEQUENCING EFFORT

Complete-MAG equivalents (equation 1) as a function of sequencing effort was modeled for environmental sequence datasets using the Gompertz equation, as reformulated by Zweitering et al. [19] for use with microbial growth curves:

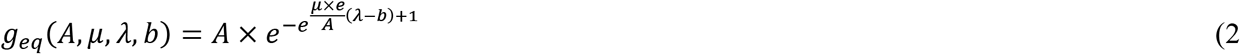

where *A*, *μ*, and *λ* are fit coefficients and *b* is high-quality bases (Gbp). From equation 2, MAG yield could be defined as:

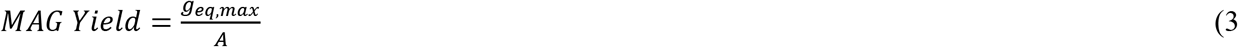

where *g_eq,max_* is the largest complete-MAG equivalent derived from the subsampling experiment and *A* is from equation 2.

### DEFINING THE MICROBIAL METAGENOME AND SEQUENCING PROBABILITY

Here we draw on set theory to provide a theoretical grounding for our *in silico* simulations described below. The expected number of sequences to sequence a fraction of an individual microbe’s genome can be modeled with probability theory by defining a community metagenome with set theory. Fig. 8a-e provides a cartoon illustrating the application of this set theory on a hypothetical microbial population, *G*. *G* contains unique genomes (*g*) with finite abundances (*n*). The definition of microbial species is somewhat contentious [32]. Here we define *g* as the average genome (length and composition) for all individuals genomes associated with the rank, species, based on an arbitrary definition. As such, the species richness (*s* or unique *g*) of *G* depends on the definition of *g*. In the example community (Fig. 8a-e), *s*=6 and *n*=13. Thus, *G* is represented as (Fig. 8a):

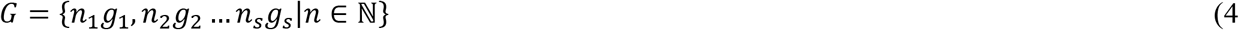

where *s* is the species richness. When characterizing *G* via shotgun metagenomics, the *i*^th^ genome, *g*_i_, can be sequenced at *K* unique sections given a characteristic read length, *k*, and average genome size, *l*, in number of base pairs (Fig. 8b). Thus, the number of physically-distinct *k*-sized reads, *K*, associated with the *i*^th^ genome, *g*_i_, within *G* is equal to:

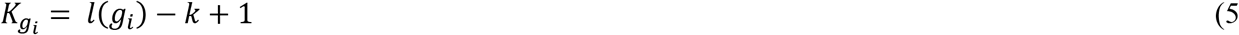

**Fig. 8.**
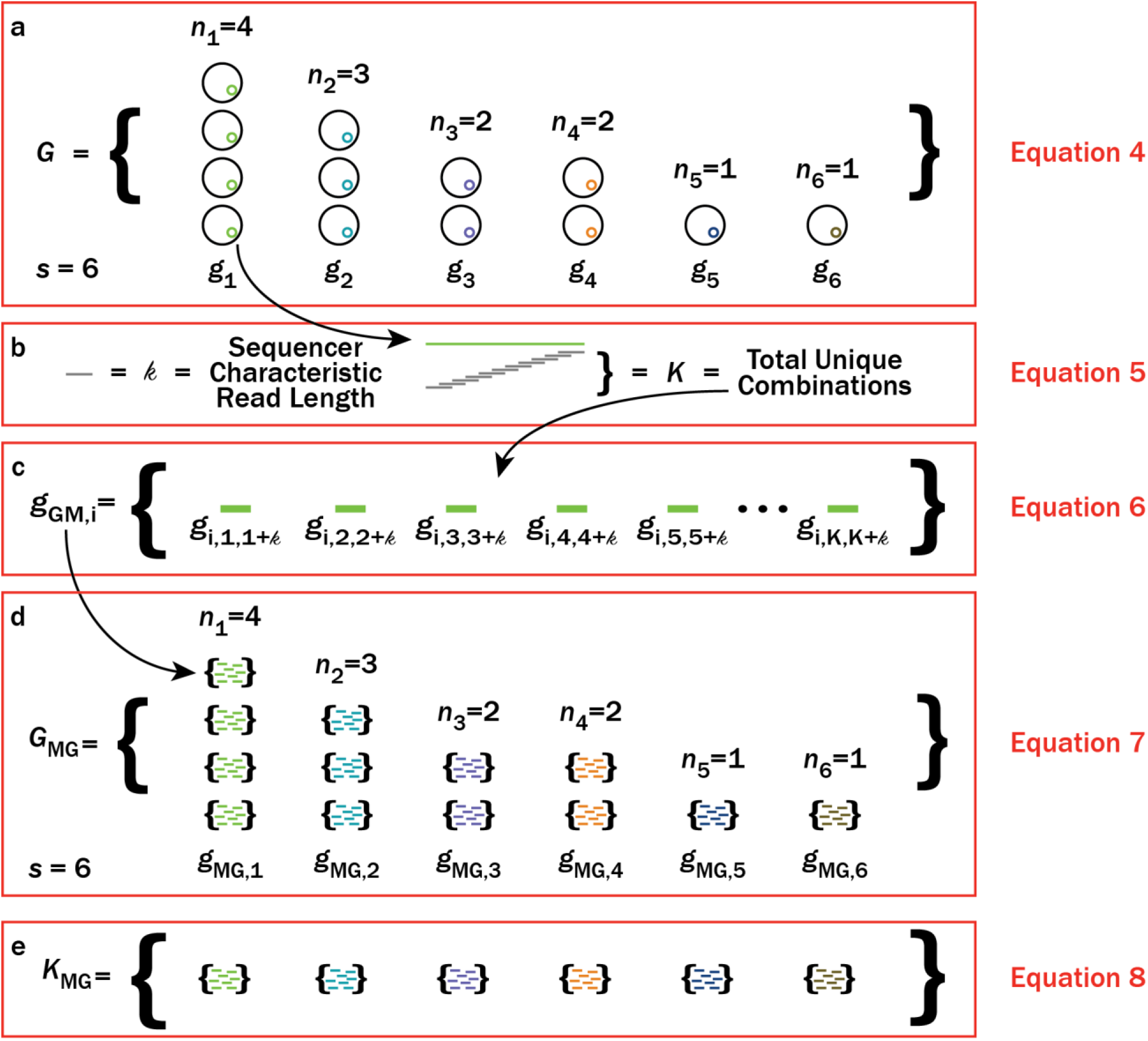
A cartoon illustrating an example microbial community (*G*), the metagenomes for genomes (*g*_MG,i_) (as defined in equation 5) within *G*, and the overall metagenome for the given microbial community (*G*_MG_). In this example, there are 6 genomes (*s*=6) and a total of 13 individual microbes. (a) Black circles represent individual microbes whose genomes are averaged together, *g*. The average genome, *g*, are indicated by different color inner-circles. (b) Individual average genomes can be sequenced at *K* unique positions depending on the characteristic read length, *k*, of a sequencer. (c) All unique positions that can be sequenced for a given genome, *g*, defines the metagenome, *g*_MG_, for the *i*^th^ genome, *g_i_*. (d) Replacing all individual genomes in (a) with metagenomes, *g*_MG_, gives the metagenome of the microbial community, *G*_MG_.

Note that equation 5 considers homologous DNA as unique fragments. From equation 5, the metagenome, *g*_MG_, for *g*_i_ is defined as the set of all physically-distinct *k*-sized reads (Fig. 8c) or:

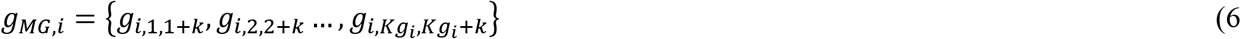

where the subscripts for *g*_i_ represent a given *k*-sized read spanning from an arbitrary starting base pair to the arbitrary starting base pair plus *k*. By substituting *g*_MG,i_ into all *g* for equation 4 (Fig. 8d), the metagenome for a microbial community, *G*_MG,_ is derived to be:

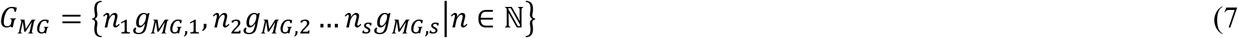

while the population of physically-distinct *k*-sized reads in the metagenome, *G*_MG_ (Fig. 8e), is represented as:

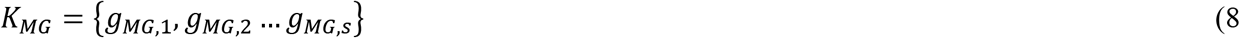

From equation 8, one can determine the cardinality, or the total number, of physically-distinct *k*-sized reads associated with *G*_MG_ (expressed as |*K_MG_*|). To an effect, |*K_MG_*| is analogous with “metagenomic richness” of an environment. When attempting to fully sequence *G*_MG_ using shotgun metagenomics, we assume that sampling events (sequence reads) are independent and are sampled with replacement. In fact, Illumina sequencing technology sequences reads in parallel via the individual DNA fragments binding to individual clusters. Furthermore, the fragmented DNA cannot be sequenced twice as the sequencing process is destructive [33]. Nonetheless, the mass of DNA extracted from a target environment represents a negligible fraction of the total DNA existing in that environment. As the relative abundance of the *k*-sized reads in *K_MG_* does not change when DNA is extracted from an environment, sampling events can be treated as independent and thus, DNA sampling reduces to sampling with replacement. If the proportion DNA mass extracted had a significant impact on the remaining mass of DNA in the environment, then one would be more suited to sequence all the DNA versus a smaller proportion of the DNA. The sequencer should have no impact on sampling assuming no sequencing errors due to misreading or spatial sampling issues (i.e., clonal density issues). These issues do exist, but for the sake of a first order approximation, these biases can be ignored.

The probability of sequencing all elements in *G*_MG_ reduces to a coupon collectors problem [34] by making the above assumptions. Using the general functional form for calculating expected samples for sampling all unique elements in a set (equation 13b in [8]), one can predict the number of sequences necessary to sequence all elements in *K*_MG_, such that the expected number of sequences, E(*G_MG_*), is:

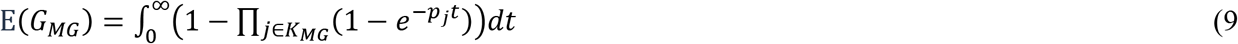

where *j* is a given element within *K*_MG_, *t* is the number of sampling events, and *p*_j_ is equal to the proportion of the *j*^th^ *k*-sized read within a given population of *k*-sized reads. *p*_j_ can be expressed as follows:

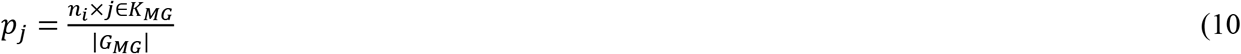

where *n*_i_ is the respective abundance for the species whose MAG contains the *j*^th^ *k*-sized read within *K*_MG_, and |*G_MG_*| is the cardinality of *G*_MG_, or the total number of *k*-sized reads in the metagenome, *G*_MG_.

### MODELING EXPECTED SEQUENCES

Equation 9 provides an estimate for the total number of sequences to sequence all *K*_MG_. The influence of increasing species richness (i.e., *s* in equation 4) on the expected number of sequences was tested for four hypothetical communities. The first community had an even structure such that all physically-distinct DNA fragments were equally distributed across all *K*_MG_. In the second community, 90% of the physically-distinct DNA fragments were equally distributed in 50% of *K*_MG_, and the remaining 10% of the physically-distinct DNA fragments were distributed equally across the remaining 50% of *K*_MG_. This community represented a community with relatively moderate species evenness. In the third community, 90% of the physically-distinct DNA fragments were equally distributed across 10% of *K*_MG_, and the remaining 10% of the physically-distinct DNA fragments were distributed equally across the remaining 90% of *K*_MG_. This community represented a community with relatively low species evenness. The last community had 10 equally-sized groups. The abundance of the physically-distinct DNA fragments in each group was based on the function form of a lognormal community [35] which has been observed in microbial populations (e.g., [17,36]), such that:

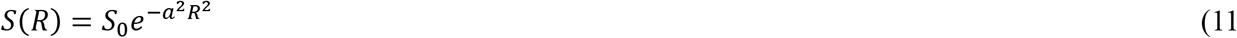

where *S*_0_ was treated as the maximum relative of abundance (*S*_0_ = 1), *a* was the inverse width of the distribution, *R* was treated as the positive octave range spanning 0 to 9, and *S*(*R*) represented the abundance for a given group. For the lognormal abundance distribution in Fig. 3d, *a* was set to a value of 0.2. Each hypothetical community started with |*K_MG_*| = 1 × 10^6^. |*K_MG_*| incrementally increased at 10 equally-spaced, linear steps to a maximum of |*K_MG_*| = 1 × 10^8^. As |*K_MG_*| increased, all community structures remained constant. Graphical representation of rank abundance in Fig. 3a-d was normalized by a given |*K_MG_*| to reflect that populations retained the same structure even as population size varied. We defined a normalized rank abundance *r*_n_ such that

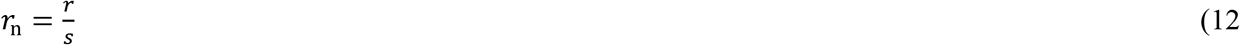

where *r* and *s* are untransformed rank abundance and richness, respectively. For each community, at each step, the expected number of sequences was calculated using equation 9. The expected number of sequences as a function of |*K_MG_*| were modeled with linear regressions.

Equation 9 gives the expected number of sequences required to sequence any sized community to exhaustion. Numerical sequencing simulations were performed to determine the number of sequences necessary to sequence a subset of all physically-distinct DNA (*K*_MG_). These numerical sequencing simulations were applied to four hypothetical community structures described above. Numerical simulations were performed such that |*K_MG_*| = 3 × 10^7^, 4 × 10^7^, 5 × 10^7^, 7 × 10^7^, 9 × 10^7^, and 1 × 10^8^. During each of these simulations, the parameters read length (*k*) and average genome size (*l*) were set to 100 and 1 × 10^6^, respectively, for all *g*. Random elements from *K*_MG_ were selected with replacement to simulate sequencing events. Numerical simulations were performed until the percentage of |*K_MG_*| sequenced was 50%, 70%, 90%, 95%, 99%, or 100%. A weight distribution was applied to elements in a given *K*_MG_. The weight distribution biased sequencing to reflect the relative abundances of the four hypothetical communities described above. The percentage of |*K_MG_*| sequenced was evaluated every 1 × 10^7^ sequences. Numerical simulations were performed in triplicate for all |*K_MG_*| and all target fractions of |*K_MG_*|.

We explored the influence of community evenness on required sequencing effort by performing sequencing simulations on 6 different lognormally-distributed communities. The numerical sequencing simulations followed the simulations described above. The 6 lognormal communities were modeled such that each community had *S*_0_ =1, *R=*10, and |*K_MG_*| = 1 × 10^7^. Values of *a* for the 6 lognormal distributions were *a*=0, *a*=0.005, *a*=0.008, *a*=0.01, *a*=0.015, and *a*=0.02. Evenness was represented using the Pielou evenness index [11], or the ratio of the Shannon diversity index [37] for an observed community to an even community of equal richness. Shannon diversity was calculated in the context of a metagenomes such that:

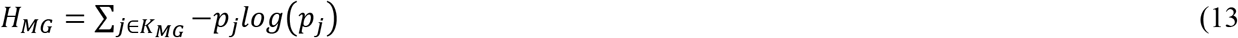

where *p*_j_ is the proportion that the *j*^th^ *k*-sized read represents among all physically-distinct DNA sequences in the metagenome. Thus, the Pielou evenness index [11] was calculated such that:

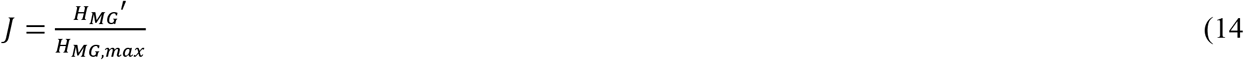

where *J* was the Pielou evenness index, *H_MG_*′ was the metagenome Shannon diversity index, and *K_MG_*_,*max*_ represented the metagenome Shannon diversity index when all *p*_j_ were equal (*a*=0).

Lastly, numerical simulations were performed to determine the sequencing effort necessary to achieve a target fraction for an individual metagenome (*g*_MG_). Target fractions increased from 0.5 to 1 at 100 linearly-spaced intervals. The fraction of the metagenome community (*G*_MG_) that *g*_MG_ represented varied from 1% to 100% in 30 lognormally-spaced intervals. The target genome sizes varied such that *l*=0.5×10^6^, *l*=1×10^6^, *l*=2×10^6^, *l*=3×10^6^, *l*=5×10^6^, *l*=10×10^6^, *l*=15×10^6^, and *l*=20×10^6^. The sequencing effort for a given combination of target fraction, genome size, and fraction of the metagenome community was modeled using the GAM function (mgcv R package; [38]). For modeling purposes, target fraction was raised to the 6^th^ power and both genome size and sequences were log-transformed. The number of smooth dimensions for fraction of community, genome size, and target fraction were heuristically varied till the resulting fit demonstrated residuals with a normal distribution. Note that the objective here was not build a predictive model but simply a first order approximation for simulations performed here. The GAM was extrapolated for genome relative abundances spanning 0.01 to 0.0001 at all sizes and target fractions. The extrapolation was necessary due to the computational limitation of analyzing genome relative abundances <0.01.

### RELATING MAG RESPONSE TO THE THEORETICAL SEQUENCING MODEL

Sequencing effort (in base pairs) used for predicting complete-MAG equivalents for the sequence read datasets maize soil, an estuarine sediment, the surface ocean, and the human gut, were related to a genome relative abundance utilizing the GAM presented in the previous section. This was accomplished by setting a constant genome size and target fraction (0.5) and performing a linear regression between genome relative abundance (a genome’s fraction of a community) and log-transformed base pairs. The linear regression was performed with genome sizes of 1 Mbp, 5 Mbp, and 20 Mbp. Complete-MAG equivalents were predicted as a function of genome relative abundance sequenced to a target fraction of 0.5.

## Declarations

### Ethics approval and consent to participate

Not applicable.

### Consent for publication

Not applicable.

### Authors’ Contributions

TMR and ADS both conceived the study, performed data analyzed, interpreted results and wrote the manuscript.

### Data Availability

All simulations and codes used for modeling sequencing effort are freely available on Github at: https://github.com/taylorroyalty/sequence_simulation_code. All data generated during the subsampling experiment is available in Additional file 1. The GRASE GUI application is available at: http://adsteen.shinyapps.io/grase.

## Acknowledgements

We also would like to thank Zachery Stooksbury for last minute assistance with editing figures.

